# *De novo* assembling a high-quality genome sequence of Amur grape (*Vitis amurensis* Rupr.) gives insight into *Vitis* divergence and sex determination

**DOI:** 10.1101/2023.10.09.561595

**Authors:** Pengfei Wang, Fanbo Meng, Yiming Yang, Qian Mu, Tingting Ding, Huiping Liu, Fengxia Wang, Ao Li, Qingtian Zhang, Shutian Fan, Bo Li, Zhiyao Ma, Tianhao Zhang, Yongfeng Zhou, Hongjun Zhao, Xiyin Wang

## Abstract

To date, there is no high-quality sequence for genomes of the East Asian grape species, hindering biological and breeding research efforts to improve grape cultivars. This study presents a ∼522 Mb of the *Vitis amurensis* (*Va*) genome sequence containing 27,635 coding genes. Phylogenetic analysis indicated that *V. riparia* (Vr) may firstly split from the other two species, *Va*, *V. Vinifera* (*Vv*; Pinot Noir: PN40024 and Cabernet Sauvignon). Much divergent gene reservation among three grape duplicated gene sets suggests that the core eudicot common hexaploidy (ECH), 130 million years ago (Mya), has still played a non-negligible role in grape species divergence and biological innovation. Prominent accumulation of sequence variants might have improved cold resistance in *Va*, resulting in a more robust cold resistance gene regulatory network than those in *Vv* and *Vr*. In contrast, *Va* preserved much fewer NBS disease resistance genes than the other grapes. Notably, multi-omics analysis identified one trans-cinnamate 4-monooxygenase gene positively correlated to the resveratrol accumulated during *Va* berry development. A selective sweep analysis revealed a hypothetical *Va* sex-determination region (SDR). Besides, a PPR-containing protein-coding gene in the hypothetical SDR may be related with sex determination in *Va*. The content and arrangement order of genes in the putative SDR of female *Va* were similar to the SDR of female *Vv*. However, the putative SDR of female *Va* lost one Flavin-containing monooxygenases (FMO) and contained one extra uncharacterized protein-coding gene. These findings will improve the understanding of *Vitis* biology and contribute to the improvement of grape breeding.

## Introduction

Grape genus (*Vitis*) contain about 60 species of vining plants. Eurasian grape, East Asian grape, American grape, and other intergeneric hybrid grapes are some of the most common grape varieties globally (Foria *et al*., 2022).

As a wild plant, originated in eastern Asia, Amur grape (*Vitis amurensis*), also called “Chinese wild grape” or “Shanputao” (Wang *et al*., 2019; Wang *et al*., 2021), is one of the East Asian grape species, resembling the wine grape (one of the Eurasian grapes). Wild Amur grapes are mainly found in the Yellow River basin and Songhua River basin in China, Siberia in Russia, Korea, Japan, and many other areas around the world (Chen *et al*., 2018). Amur grapes have female, male, or bisexual plants (Figure 1b-d). The Latin name “*Vitis amurensis* Rupr.” of the Amur grape was first proposed in 1857 by botanist Franz Josef Ruprecht, and the cultivation of *Vitis amurensis* can be traced back to 1907 (wine-world.com). A female Amur grape cultivated variant (cv), called “Zuoshan No. 1” is vastly planted in the Northern China. A bisexual Amur grape cv. “Shuangyou” is also widely cultivated.

**Figure 1.**
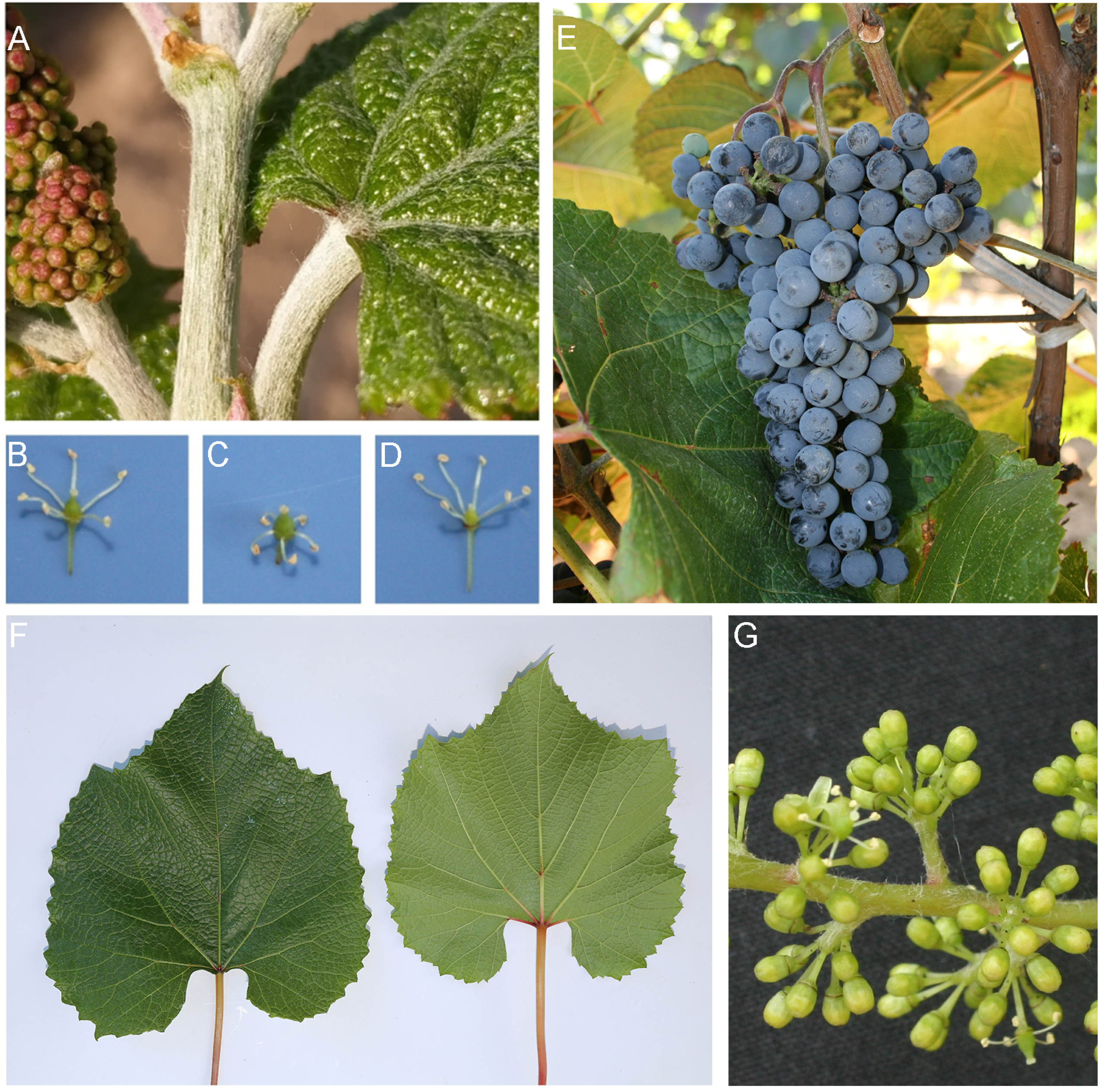
Amur grape Atlas. (A) Spider silk-like hairs of Amur grape. (B) The bisexual flower. (C) The female flower. (D) The male flower. (E) Amur grape cv. Zuoshan No.1. (F) The leaf of Zuoshan No.1. (G) The flower of Zuoshan No.1.

Amur grape has huge commercial potential, for containing abundant phenolic compounds, procyanidins, oligostilbenes, and stilbenes (Chen *et al*., 2018). Its fruits are dark purple with small berries, more acidic than *Vv*, and are used as raw materials for claret, sweet red wine, semi-dry wine, semisweet red wine, and brandy (wine-world.com). Young twigs of the Amur grape have spider silk-like hairs (Figure 1a). Amur grape is extremely cold-tolerant and can survive temperatures as low as -40°C (Wan *et al*. 2008; Xu *et al*. 2014). Now, Amur grape has become a valuable germplasm resource for grape breeding and wine production. Cv. “Beibinghong”, a cross of *Vitis amurensis* and *Vitis vinifera,* is a good example in cold resistance, having ability to tolerate temperatures of -37 °C and therefore used for preparing fancy red ice wine. Recent pharmacological studies showed that Amur grape possesses anti-inflammatory, antimicrobial, antioxidant, and anticancer properties (Chen *et al*., 2018).

The genome sequence of *Vitis Vinifera* cv. “Pinot Noir (PN40024)”, a kind of Eurasian grape species, was deciphered (Jaillon *et al*., 2007; Canaguier *et al*., 2017) and has been used as the standard grape genome for grapes’ gene analyses (Liang *et al*., 2019; Ma *et al*. 2018). A high-quality genome sequence of *Vv* cv. Cabernet Sauvignon was sequenced in 2021 with the Hi-C technology. The genome for *V. riparia*, an American grape species, was recently decoded (Girolletet al., 2019). *Va* IBCAS1988 genome was sequenced in 2021 with a genome size 604.56 Mb and N50 282,256 bp (Wang *et al*., 2021). To date, there is no high-quality sequence for genomes of the East Asian grape species, hindering biological and breeding research efforts to improve grape cultivars.

Here, we present a high-quality sequence for the *Vitis amurensis* cv. “Zuoshan No.1” genome. The availability of the genome sequence of Amur grape will lay a solid foundation to understand grape biology and develop new breeding lines, by deepening our insight into grapes’ evolution and genetic variability, the molecular basis for cold adaptation and sex determination of grapes, etc.

## Results

### A high-quality Amur grape reference genome

The study used the female Amur grape “Zuoshan No.1” (Figure 1E-G) as the material. The Amur grape genome sequence was *de novo* assembled with 52.69 Gb (∼101.46 fold coverage) of Nanopore reads, 98.79 Gb (∼189.23 fold coverage) of MGISEQ short reads, and 58.12 Gb (∼111.34 fold coverage) of Hi-C data. A k-mer analysis of Illumina reads estimated the Amur grape genome to be ∼532 Mb, with DNA heterozygosity 1.20% (Table 1). The final assembled genome sequence reached to ∼522 Mb, covering ∼97.5% of the estimated genome (BUSCO value = 97.5), with a GC content 0.345 (Table 1). Moreover, the assembly comprised 56 contigs with 2.5 Mb of contig N50. The N50 of *Va* IBCAS1988 is much shorter (282,256 bp) (Wang *et al*., 2021) than that of the *Va* Zuoshan No.1 genome, suggesting quality differences in the genome assemblies. The resultant 615 contigs accounted for 98.31% (∼513.47Mb) of the total assembled genome, anchored onto 19 pseudo-chromosomes using Hi-C reads (Figure 2A and B). Repetitive elements accounts for 59.21% of the genome sequence (Table 1), much higher than the finding (47.06%) in *Va* IBCAS1988 genome (Wang *et al*., 2021).

**Figure 2.**
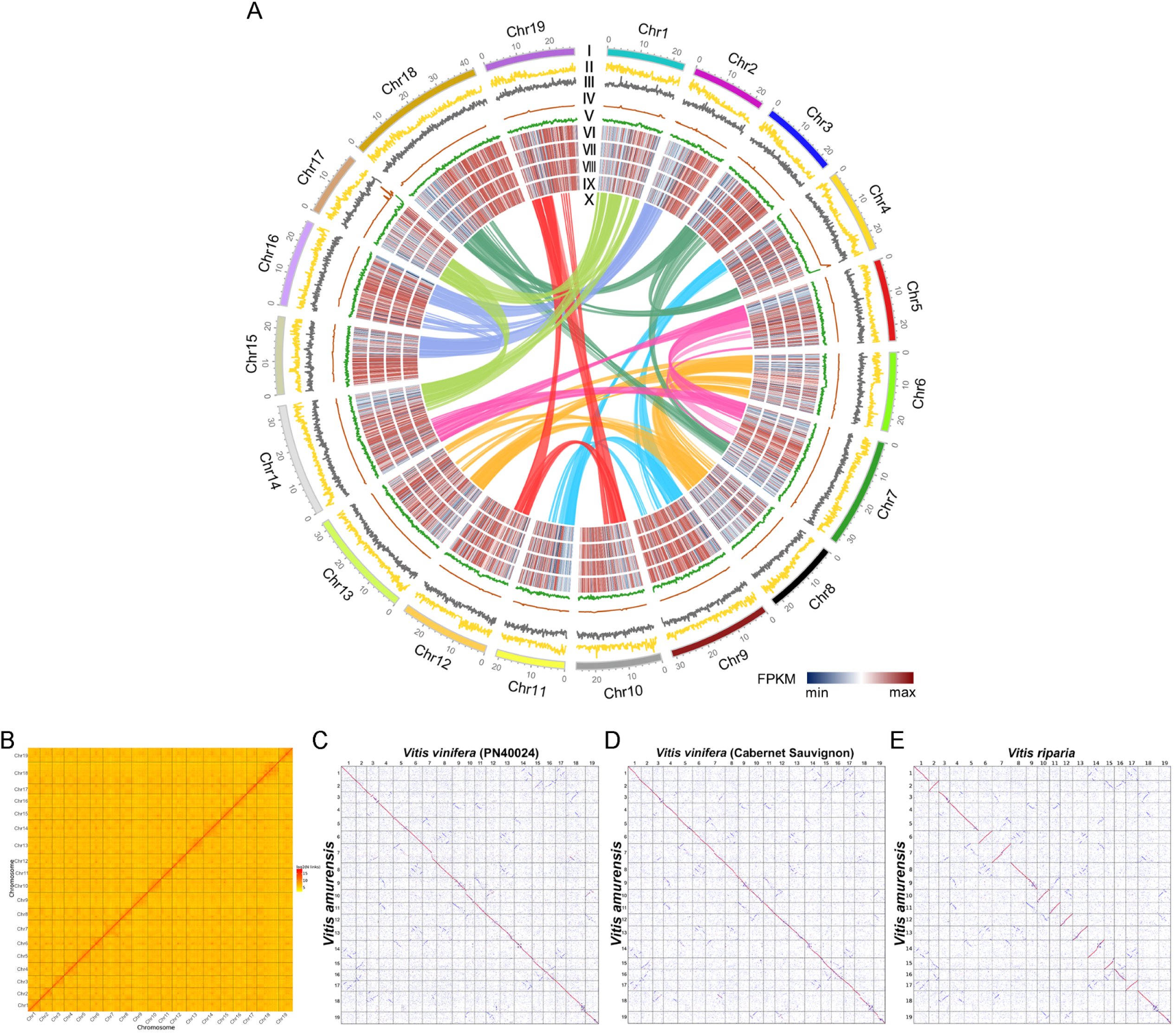
Genomic feature of *Va* Zuoshan No.1 and comparison with other genomes. (A) The landscape of genome assembly and annotation of *Va* Zuoshan No.1. Tracks from outside to the inner correspond to I, chromosomes; II, gene density; III, repeat density IV, Non-coding RNA density; V, GC content; VI-IX, Gene expression levels (FPKM) in S4-S1; and X, synteny information. (B) Hi-C map of *Va* Zuoshan No.1. (C) Dotplot of homologous genes between *Va* Zuoshan No.1 and *Vv* PN40024. (D) Dotplot of homologous genes between *Va* Zuoshan No.1 and *Vv* Cabernet Sauvignon. (E) Dotplot of homologous genes between *Vv* Cabernet Sauvignon and *Vr*.

**Table 1.** Summary statistics for the *Va* Zuoshan No.1 genome assembly in comparison with the *Va* IBCAS1988 genome.

| Genomic feature | Zuoshan No.1 | IBCAS1988 |
| --- | --- | --- |
| Estimated genome size | 532.35Mb | 607Mb |
| Total assembly size | 522.28Mb | 604.56Mb |
| Number of contigs | 615 | — |
| Largest contig | 14,947,757bp | 2,623,866bp |
| Contig N50 length | 2,512,611bp | 282,256bp |
| Scaffold N50 | 26,519,999bp | 748,673bp |
| Sequences anchored to chromosomes | 513.47 Mb/98.31% | — |
| GC content | 34.50% | 34.37% |
| Complete BUSCOs | 97.50% | — |
| Repeat(%) | 59.21% | 47.06% |
| Protein-coding genes | 27,635 | 32,885 |
| Heterozygous ratio(%) | 1.20% | 1.23% |

We predicted 27,635 protein-coding genes using an integrative strategy combining *de novo* gene prediction, protein-based homology search, and transcript data from RNA sequences of various tissues. The ∼96.4% of annotated protein-coding genes could be annotated by at least one public database (InterPro, Nr, GO, KOG, Swissprot, TrEMBL and KEGG) (Table S1). Moreover, 231 miRNAs, 651 tRNAs, 291 rRNAs, and 412 snRNAs were annotated. LTR accounts for the largest proportion of transposable elements, making up ∼46.39% of the genome.

The homologous gene dotplotting analysis showed a big difference in arrangement order of homologous genes between the *Va* Zuoshan No.1 and *Va* IBCAS1988 genomes (Figure S1). Moreover, homologous gene dotplots and chromosome synteny analysis showed highly similar arrangement order of homologous genes across *Va* Zuoshan No.1, *Vv* PN40024, *Vv* Cabernet Sauvignon, and *Vr* genomes (Figure 2C-E). *Va* has large inversions on chromosomes 3, 9, 10, and 18 relative to *Vv* and *Vr* (Figure 2D, E). Besides, *Va* IBCAS1988 chromosomes have large inversions on *Va* Zuoshan No.1 chromosomes 1, 5, 15, 16, 18, and 19 (Figure S1). The homologous gene dotplots also showed a major difference between *Va* IBCAS1988 and *Va* Zuoshan No.1 genomes on chromosome 13. The homologous region between chromosome 13 of *Va* IBCAS1988 and *Va* Zuoshan No.1 is clearly divided into two parts. The anterior part of chromosome 13 (∼19.6 Mb) of *Va* IBCAS1988 is homologous to the posterior part of *Va* Zuoshan No.1 chromosome while the posterior part of *Va* IBCAS1988 chromosome is homologous to the anterior part of *Va* Zuoshan No.1 chromosome (∼12.6 Mb) (Figure S1). Previous studies on the *Va* IBCAS1988 genome indicated a major difference between *Va* IBCAS1988 and *Vv* PN40024 on chromosome 13 (Wang *et al*., 2021). Here, we found a minimal difference between *Va* Zuoshan No.1 and *Vv* PN40024 on chromosome 13 (Figure 2C).

In order to discover the differences between the previous genome and our genome. The “putative structural variation (SV) analysis” of genomes based on the genome alignment showed big differences in the genome structures between *Va* IBCAS1988 and *Va* Zuoshan No.1 genomes. This study identified 171 inversions (95.46Mb), 794 translocations (9.86 Mb), and 481 duplications (3.41Mb) (Table S2) between the Amur grape Zuoshan No.1 and *Va* IBCAS1988 genomes. The SV analysis of the genome alignment proved the large inversions on chromosomes 1, 5, 15, 16, 18, and 19 shown by the homologous gene dot plots. Moreover, the SV analysis also indicated large inversions on chromosomes 3, 8, 9, 10, 11, 14, and 17. However, the 122 duplications between the Amur grape Zuoshan No.1 and *Va* IBCAS1988 genomes involved 97 protein-coding genes. The 99 translocations between the Amur grape and *Va* IBCAS1988 genomes involved 82 protein-coding genes.

As to the SNPs, a total of 1,382,913 SNPs were identified between two *Va* cultivars, Zuoshan No.1 and IBCAS1988 (Table S2), including 10,733 protein-coding genes. Besides, a total of 379,188 indels were identified, with insertions affecting 7654 protein-coding genes, and deletions 8356 ones. The Amur grape Zuoshan No.1 genome had 12018 genomic-specific segments (17.51 Mb), and the *Va* IBCAS1988 had 8465 genomic-specific segments (12.25 Mb), including 2271 *Va*-specific PAV (presence-absence variation) and 237 *Va* IBCAS1988 specific PAV genes (Table S3). Many *Va* Zuoshan No.1-specific PAV genes include pentatricopeptide repeat-containing protein and TF genes (Table S4). Many *Va* IBCAS1988-specific PAV genes include hypothetical protein genes (Table S5).

### Comparative genomics analysis of three grape species

Duplicated genes produced by the core-eudicot-common hexaploidy (ECH) (Jaillon *et al*., 2007), common to major core eudicot, may have contributed to the divergence of the species. We found that *Vr* contains more duplicated genes (5231, 20.04%) than *Va* (4494, 16.26%) and *Vv* (3993, 12.49%) (Table S2), showing 800-1300 duplicated gene composition difference. This should have been caused by unbalanced gene losses in the ECH-produced regions.

Polyploidization was anticipated to result in genome instability, characterized by extensive DNA rearrangements and gene losses, especially in the early days after the tripling of the genome. Notably, the present finding suggest much divergent evolutionary patterns among these grapes and the ECH, as ancient as 130 million years ago (Mya), played and has still played a non-negligible role in grape species divergence and biological innovation.

The Ks distribution of duplicated genes generated by the ECH shows an up to 3.2% difference in their evolutionary rates. *Vv*, *Vr*, and *Va*, had peaks located at 1.29 (+/-0.150), 1.27 (+/-0.165), and 1.25 (+/-0.135), respectively, indicating that *Va* evolved the slowest among the three species (ANOVA, *P* Value=0.023) (Figure 3a, Supplementary Table 7). The Ks peaks derived from the orthologous gene pairs between species were located around 0.026 (Figure 3b, Table S3). Considering the proposed occurrence time of the ECH, and the Ks between paralogous generated by each grape ECH event was ∼1.25 (Figure 3a), we calculated the three species should have diverged ∼2.3-2.8 Mya based on the Ks between orthologous of every two grapes was ∼0.025-0.026 (Figure 3b).

**Figure 3.**
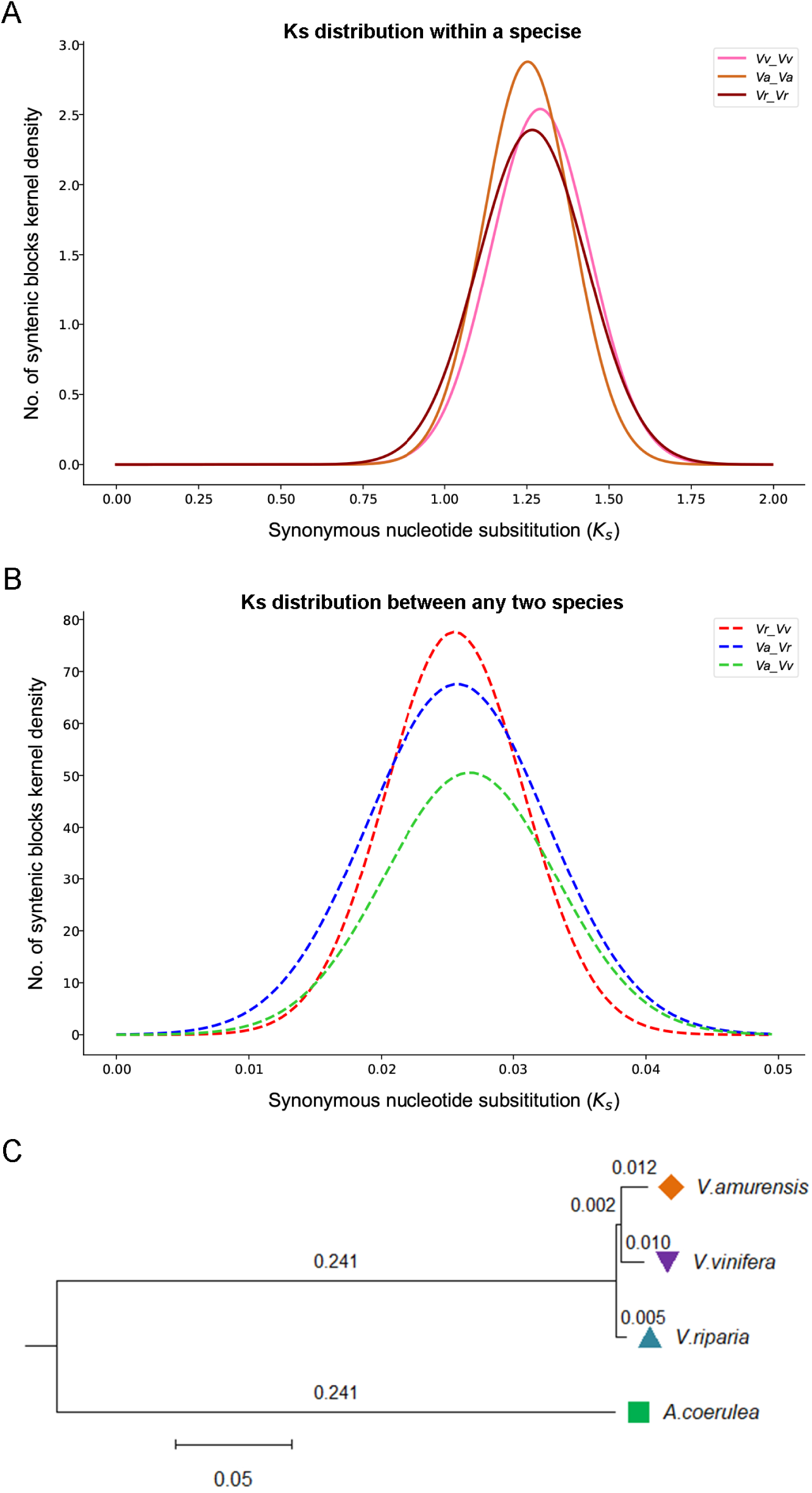
Ks distribution of collinear genes and phylogenetic relationship of species. (A) Ks distribution of homologous genes within a species. (B) Ks distribution of homologous genes between any two species. (C) Evolutionary tree of selected plants. The number is the step size and represents the size of the difference between two adjacent sequences.

By using the concatenated multiple sequence alignment of single-copy genes (5929) inferred by Orthofinder, between the three *Vitis* genomes and *Aquilegia coerulea*, acting as an outgroup, a phylogenetic tree was constructed and showed that the *Va* was more closer to *Vv* in evolution history (Figure 3C). When Arabidopsis or Boxwood was used as an outgroup, we obtained the same inference (Figure S2).

The SVs analysis revealed large DNA inversions on *Va* chromosomes 3, 9, 10, and 18, relative to *Vv* (both Cabernet Sauvignon and PN40024) and *Vr*, consistent with the homologous gene dotplots (Figure 2; Figure 4; Figure S3). Large DNA rearrangements contributed to the divergence of different grapes. DNA rearrangements in *Vitis* chromosomes 19 were inferred across the three species, large inversions were found between the *Vr* and *Vv* chromosomes 3, 7, and 18, especially one large DNA segmental translocation was detected on *Vr* and *Vv* chromosome 18 (Figure 4; Figure S3).

**Figure 4.**
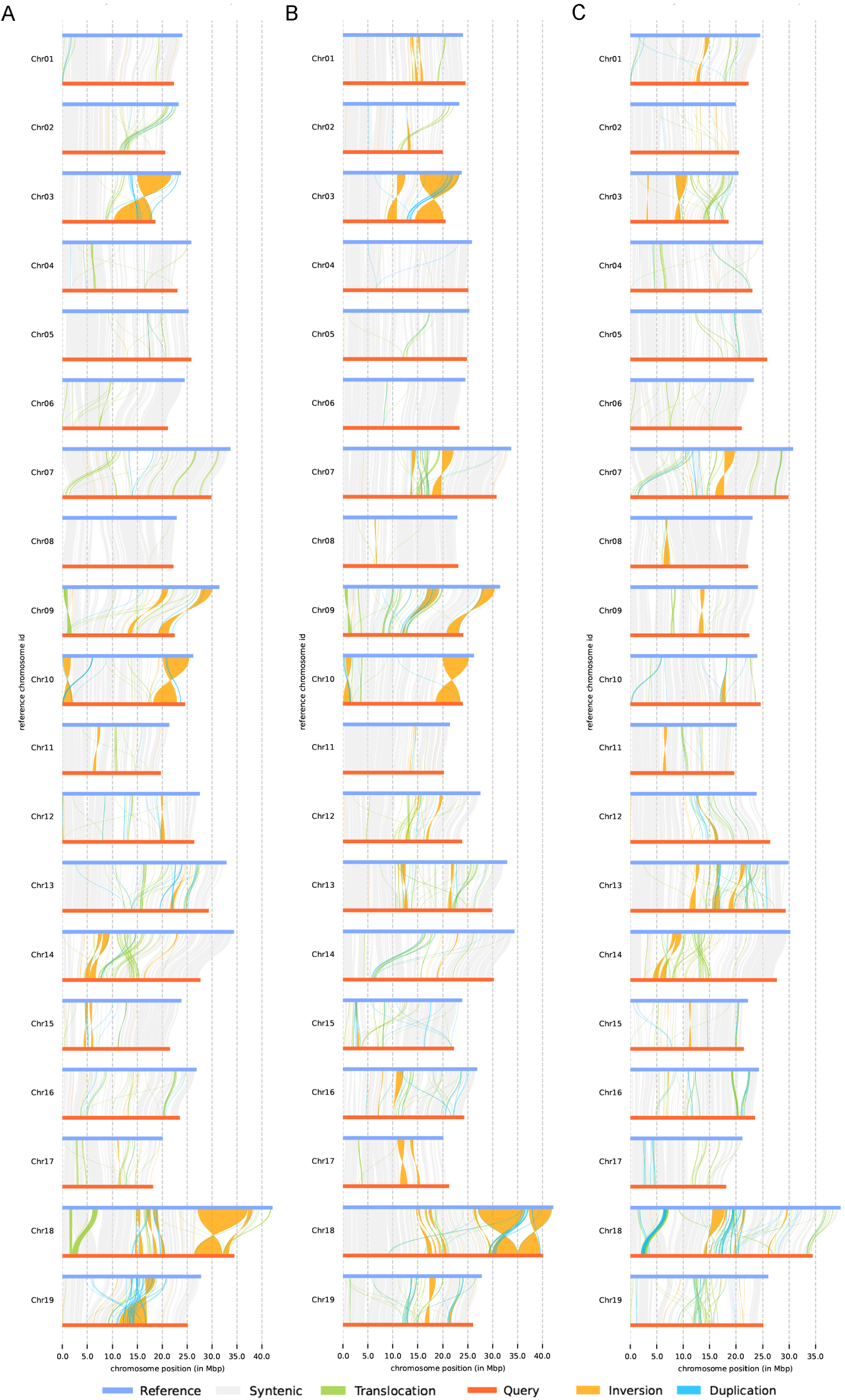
Structural Variation analysis. (A) A comparison between *Va* and *Vv*, with *Va* as the reference. (B) A comparison between *Va* and *Vr*, with *Va* as the reference. (C) A comparison between *Vr* and *Vv*, with *Vr* as the reference.

Genome instability due to the ECH resulted in species-specific regions, further contributing to their biological divergence. A comparison of *Va* and *Vv* showed that Amur grape had 12105 specific genomic segments, accumulated to 15.46 Mb, whereas *Vv* had 8465 (11.73 Mb), with 1609 *Va*-specific PAV genes and 79 *Vv*-specific PAV genes positioned within these specific genomic segments (Table S4). The *Va*-specific PAV genes include those respectively encoding cold-responsive protein kinase 1, ethylene-responsive transcription factor, flavonoid 3’,5’-hydroxylase 2, DELLA2, heat shock 70 kDa protein, methyl-CpG-binding domain protein, and WRKY transcription factor 14 genes (Table S5). In contrast, *Vv*-specific PAV genes contain several uncharacterized genes (Table S6). We found 16 *Va*-specific PAV genes were cold resistance related genes. The enriched GO terms of *Va*-specific PAV genes (P<0.05) comprise monolayer-surrounded lipid storage body (GO:0012511) and stress fiber (GO:0001725). The enriched KEGG pathway of *Va*-specific PAV genes (P<0.05) comprise Calcium signaling pathway (ko04020), and phosphonate and phosphinate metabolism (ko00440).

A comparison of Amur grape and *Vr* genomes found that *Va* had 7142 specific genomic segments (6.53 Mb), while *Vr* had 3489 segments (7.68 Mb), including 266 *Va*-specific (Table S7) and 88 *Vr*-specific PAV genes (Table S8). We found the Vitis03G0393, Vitis11G0466, Vitis16G0074, Vitis14G1585, and Vitis05G0085 from *Va*-specific PAV genes were cold resistance related genes. The enriched GO terms of *Va*-specific PAV genes (P<0.05) comprise negative regulation of stomatal complex development (GO:2000122), regulation of vacuolar transport (GO:1903335) and cellular water homeostasis (GO:0009992). The enriched KEGG pathway of *Va*-specific PAV genes (P<0.05) comprise Cutin, suberine, and wax biosynthesis (ko00073), and fatty acid biosynthesis (ko00061).

We also identified 11503 *Vr* specific (19.93 Mb) and 5439 *Vv* genomic-specific segments (14.26 Mb), including 1556 *Vr*-specific and 224 *Vv*-specific PAV genes positioned within these specific segments.

We identified 349 cold resistance related genes (CRGs) in *Va*, 437 in *Vv* and 404 in *Vr*, accounting 1.26%, 1.37% and 1.01% of the total number of genes in each genome, respectively. An *in silico* interaction network of key cold resistance related genes in *Va*, *Vv*, and *Vr* was constructed as previously described (Song *et al*., 2020). Though having the fewest CRG genes in *Va*, a research into the robustness of CRG regulatory networks showed that *Va* CRG regulatory network was more robust in *Va* than those in *Vv* and *Vr* (Table 2).

**Table 2.**
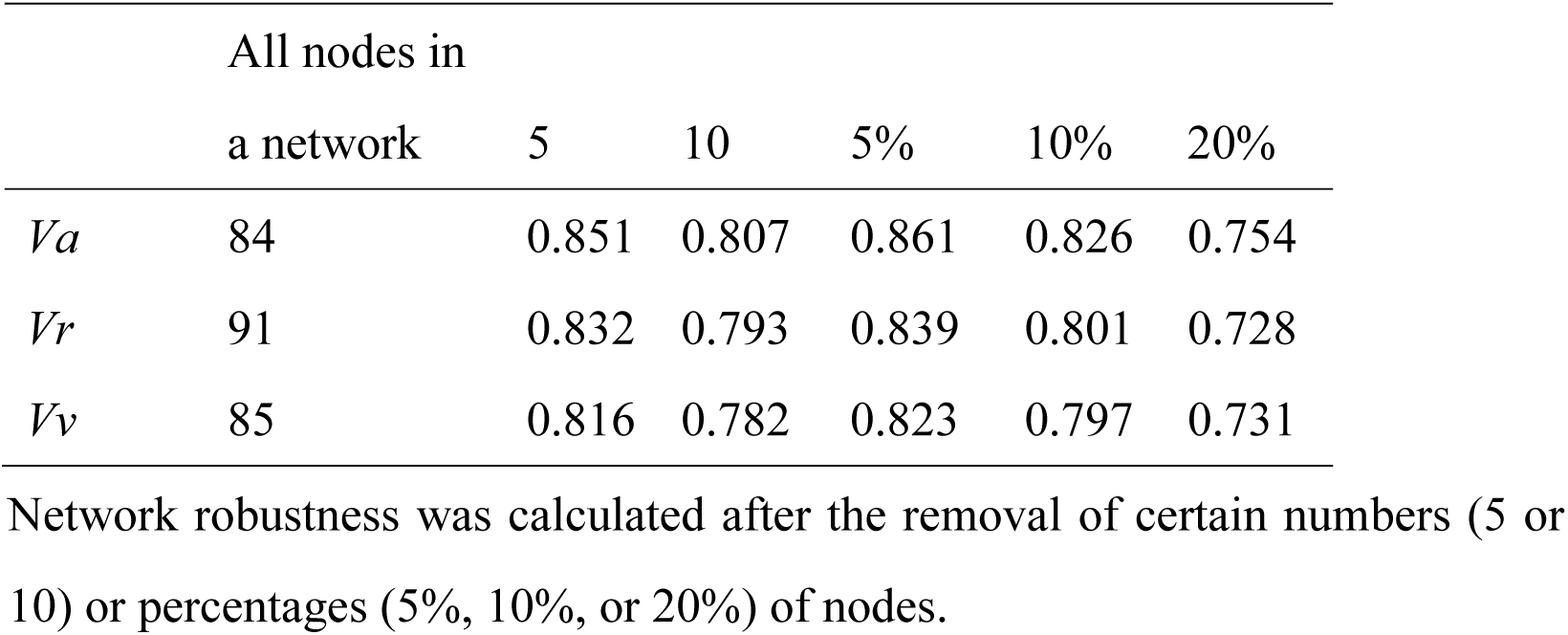
The assessment of interaction networks constructed using candidate CRGs in *Va*, *Vr* and *Vv*. Network robustness was calculated after the removal of certain numbers (5 or 10) or percentages (5%, 10%, or 20%) of nodes.

In addition to cold resistance related genes, we studied disease resistance genes. We identified totally 64 nucleotide binding site (NBS) genes in *Va*, which is less than in *Vv* (158 NBS family genes) and in *Vr* (172 NBS family genes). 34 *Va* NBS family genes (53.13% of *Va* NBS genes) were related to the ECH and 42 (65.63% of *Va* NBS family genes) were related to the tandem duplication. The Ks value of 4 pairs *Va* NBS genes is less than 0.025 (Figure S4), showing likely being generated after divergence of *Va* from the other grape species. We found 151 ancestral *vitis* NBS family genes based on the collinear relationship across three grapes and *A. coerulea* and also found that 124 of the ancestral NBS genes were lost in *Va*. In contrast, *Vv* lost 57 and *Vr* lost 54 genes. The evolutionary tree of NBS genes supports this viewpoint. Figure S5 showed in many branches which existed more than one *Vv* or *Vr* genes contain one or none *Va* gene.

### The regulatory mechanisms of nutrient accumulation at four developmental stages of *Vitis amurensis*

Berries of *Va* are nutrient-richand contain many phenolic compounds, such as anthocyanins and procyanidins, and stilbenes such as resveratrol (Chen *et al*., 2018). Besides, *Va* berries can be used in wine making because it is rich in sugars and organic acids.

To reveal the molecular mechanisms of nutrient accumulation in *Va* cv. Zuoshan No.1 berries, we profiled the RNA-seq (Table 3) and calculated the gene expression changes of key biological pathways during grape development, and correlated the gene expression changes and metabolite accumulation. Four development stages of berries, including Stage1 (S1, late period of berry expansion), Stage 2 (S2, Veraison), Stage 3 (S3, the period when berries change color completely), and Stage 4 (S4, maturity stage) (Figure 5a) were analyzed.

**Figure 5.**
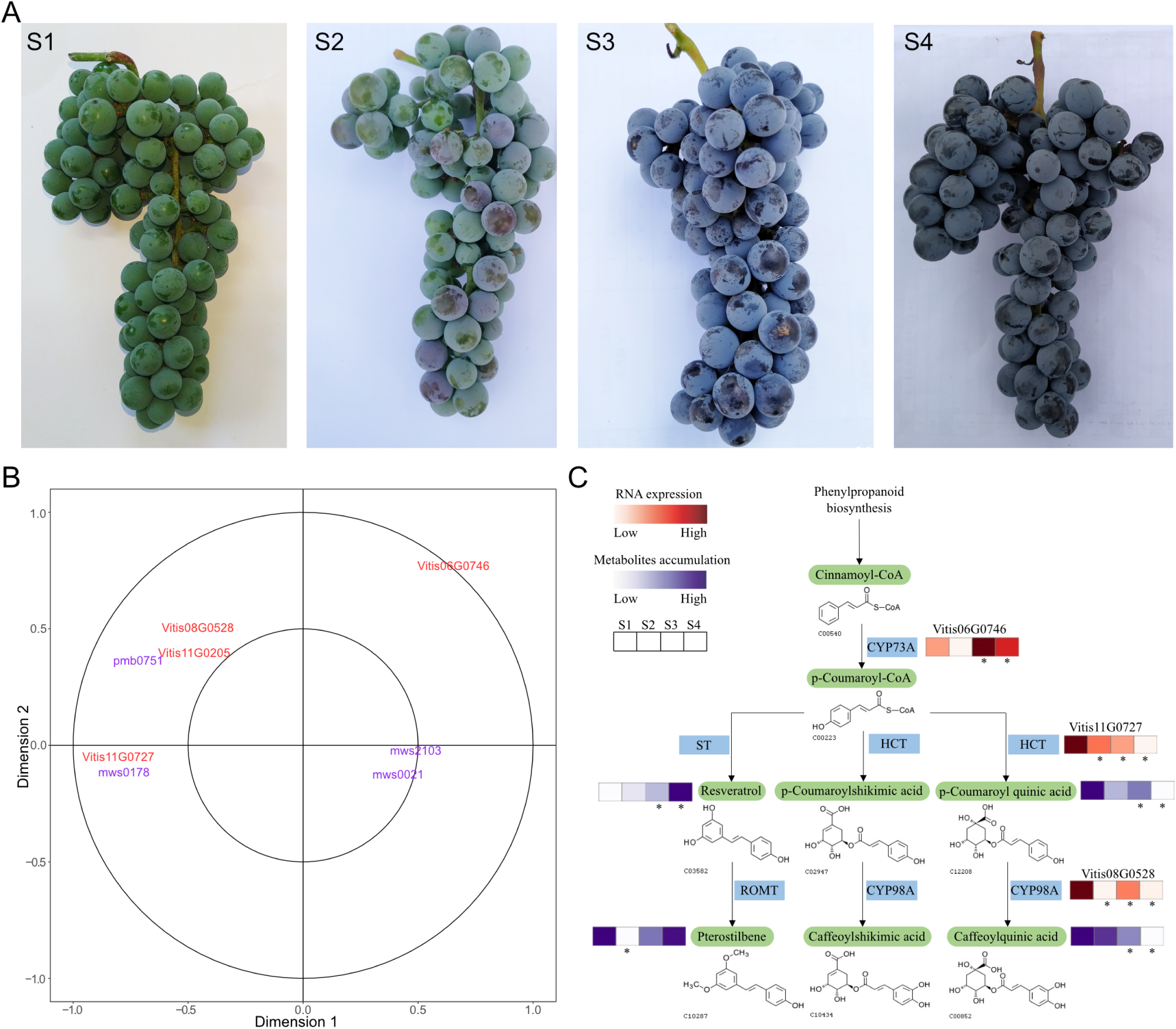
Multiomics analysis of *Vitis amurensis* fruit at different stages. (A) Fruits at stages S1, S2, S3 and S4. (B) CCA analysis of key genes and metabolites in resveratrol metabolic pathway. (C) Schematic diagram of resveratrol synthesis and metabolism mechanism during fruit ripening of *vitis amurensis*. “*” represents that these genes expression or metabolites content of the sample changes significantly compared with S1(P<0.05). CYP73A is the symbol of trans-cinnamate 4-monooxygenase, ST is the symbol of stilbene synthase, HTC is the symbol of shikimate O-hydroxycinnamoyltransferase, ROMT is the symbol of trans-resveratrol di-O-methyltransferase and CYP98A is the symbol o of 5-O-(4-coumaroyl)-D-quinate 3’-monooxygenase.

**Table 3.**
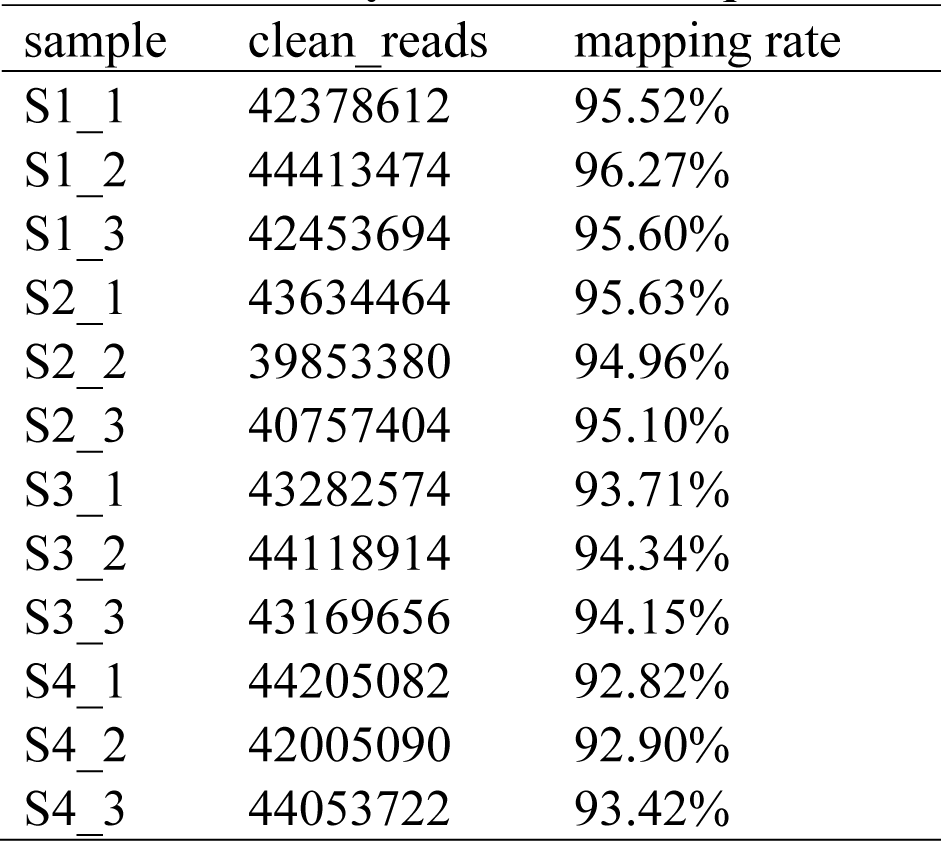
Summary of the RNA-seq.

The total sugar content increased significantly, and titratable acidity content was decreased significantly from S1 to S4 (Figure S6). Widely targeted metabolomics analysis showed that succinic acid and 2-isopropyl malic acid contents decreased significantly from S1 to S4. However, there was no significant change in tartaric acid content in berries at the four stages.

Widely targeted metabolomics analysis showed that the content of D-glucose, a monosaccharide, significantly increased between S1 and S4. Trehalose and sucrose, two disaccharides, significantly increased at S3.

Here, RNA-seq analysis is used for joint analysis with the widely targeted metabolomic analysis. Two starch and sucrose metabolism pathway (Ko00500) key genes (metabolic pathways key genes were judged based on KEGG database, and the metabolic pathways key genes were the genes could be mapped in related KEGG pathways) for sucrose synthesis (sucrose-6-phosphatase gene (Vitis08G0202)) and trehalose synthesis (alpha, alpha-trehalase gene (Vitis02G0204)) were significantly up-regulated (fold > 2, *P* value < 0.05) at S4 compared to S1. At S3, the key genes for sucrose synthesis (sucrose-phosphate synthase (Vitis05G0995 and Vitis18G2061)) and trehalose synthesis (alpha, alpha-trehalase gene (Vitis02G0204)) were significantly up-regulated. In contrast, the key gene of trehalose degradation beta-fructofuranosidase (Vitis16G0541) was significantly down-regulated (fold < 0.5, *P* value < 0.05) at S3. At S4, the key genes for sucrose synthesis (sucrose-6-phosphatase (Vitis08G0202)), sucrose-phosphate synthase (Vitis05G0995 and Vitis18G2061), trehalose synthesis (alpha, alpha-trehalase gene (Vitis02G0204)) were significantly up-regulated. In contrast, sucrose metabolism-related genes (beta-fructofuranosidase, Vitis16G0541, and Vitis02G0497) were significantly down-regulated at S4 than S2.

Widely targeted metabolomics analysis showed that the resveratrol content increased significantly from S1 to S4, but there was no significant change in the resveratrol content between berries sampled at S1 and S2. In the stilbenoid, diarylheptanoid, and gingerol biosynthesis pathway (Ko00945), stilbene synthase competitively convert the p-Coumaroyl-CoA synthesized by trans-cinnamate 4-monooxygenase to resveratrol. In contrast, shikimate O-hydroxycinnamoyltransferase and 5-O-(4-coumaroyl)-D-quinate 3’-monooxygenase could convert the same to caffeoylquinic acid. From S1 to S2, the key genes for caffeoylquinic acid synthesis, shikimate O-hydroxycinnamoyltransferase gene (Vitis11G0727) and 5-O-(4-coumaroyl)-D-quinate 3’-monooxygenase gene (Vitis08G0528) were significantly down-regulated. From S1 to S3, the trans-cinnamate 4-monooxygenase gene (Vitis06G0746) was significantly up-regulated, and the shikimate O-hydroxycinnamoyltransferase gene (Vitis11G0727) was significantly down-regulated. Meanwhile, the content of caffeoylquinic acid was significantly down-regulated, and the resveratrol content increased significantly. From S1 to S4, the trans-cinnamate 4-monooxygenase gene (Vitis06G0746) was significantly up-regulated, while the shikimate O-hydroxycinnamoyltransferase (Vitis11G0727) and 5-O-(4-coumaroyl)-D-quinate 3’-monooxygenase (Vitis08G0528) genes were significantly down-regulated. Meanwhile, the content of caffeoylquinic acid decreased significantly, while the resveratrol content increased significantly. The canonical correlation analysis (CCA) showed that the trans-cinnamate 4-monooxygenase gene (Vitis06G0746), related to the resveratrol content, and shikimate O-hydroxycinnamoyltransferase (Vitis11G0727) and 5-O-(4-coumaroyl)-D-quinate 3’-monooxygenase (Vitis08G0528) gene are also related to the caffeoylquinic acid content (Figure 5b, C).

Divergently expressed genes (DEGs) were checked (Table 4) between S1-S4. They included the putative ripening-related gene (Vitis08G0132), the chalcone synthase 2 gene (Vitis05G0886), the flavonoid 3-O-glucosyltransferase gene (Vitis16G0156), and sugar transporter SWEET2a (Vitis10G0718) gene, the ethylene-responsive transcription factor 3-like gene (Vitis12G0453), the UDP-glucose: flavonol synthase/flavanone 3-hydroxylase gene (Vitis08G0277), and some bHLH, MYB transcription factors, and WD repeat-containing protein.

**Table 4.** The number of DEGs.

| group | all | up | down |
| --- | --- | --- | --- |
| S2vsS1 | 3307 | 1367 | 1940 |
| S3vsS1 | 4633 | 1862 | 2771 |
| S4vsS1 | 5969 | 2209 | 3760 |
| S3vsS2 | 2904 | 1196 | 1708 |
| S4vsS2 | 4817 | 1904 | 2913 |
| S4vsS3 | 1797 | 652 | 1145 |

### Exploring Amur grape “putative sex-determining region (SDR)”

A previous study showed that *Vv* SDR is located on the segment between 4,801,876 bp and 5,061,548 bp on chromosome 2 and involves 15 protein-encoding genes (Massonnet *et al*., 2020). The 5’ terminus of the SDR segment has a PPR-containing protein-coding gene and the 3’terminus has an APT3 gene (Massonnet *et al*., 2020). Another study showed the SDR is located on the segment between 4,810,929 bp and 4,921,949 bp on chromosome 2 which overlapped with the previous SDR region (Badouin et al.,2020). We explored the *Vv* SDR syntenic region between 5055465 and 5198824 on Amur grape chromosome 2, showing good synteny between two species. To determine whether the Amur grape putative SDR region is related to sex, we adopted the selective sweep method, which allows analysis of fewer samples than GWAS (Kim *et al*., 2020).

Through re-sequencing and analyzing SNPs in 24 Amur grape individuals, including five male, ten female, and nine hermaphroditic individuals (including four tetraploids), we performed phylogeny analysis and examined genetic population structure among the Amur plants (Table S9). A set of 2588125 high-quality SNPs were inferred and explored to characterize the relationship among these 24 *Va* individuals and the *Va* reference genome (this study). A phylogenetic tree was constructed based on the SNPs of the 24 *Va* individuals and indicated that the individuals could be divided into two groups. However, the grouping proved unrelated to sex division (Figure 6A). Then, a genetic population structure analysis was performed and showed that the individuals could be divided into two groups, and the grouping pattern proved unrelated to sex division, either (Figure 6B). Alternatively, based on the SNPs in the SDR regions of the 24 individuals, we reconstructed a phylogenetic tree. Notably, we found that the tree had groups consistent to sex division (Figure 6C). Grossly, the above analysis showed that the genetic relationship at the population level is not related to sex division, while the SNP difference in SDR regions is related to gender.

**Figure 6.**
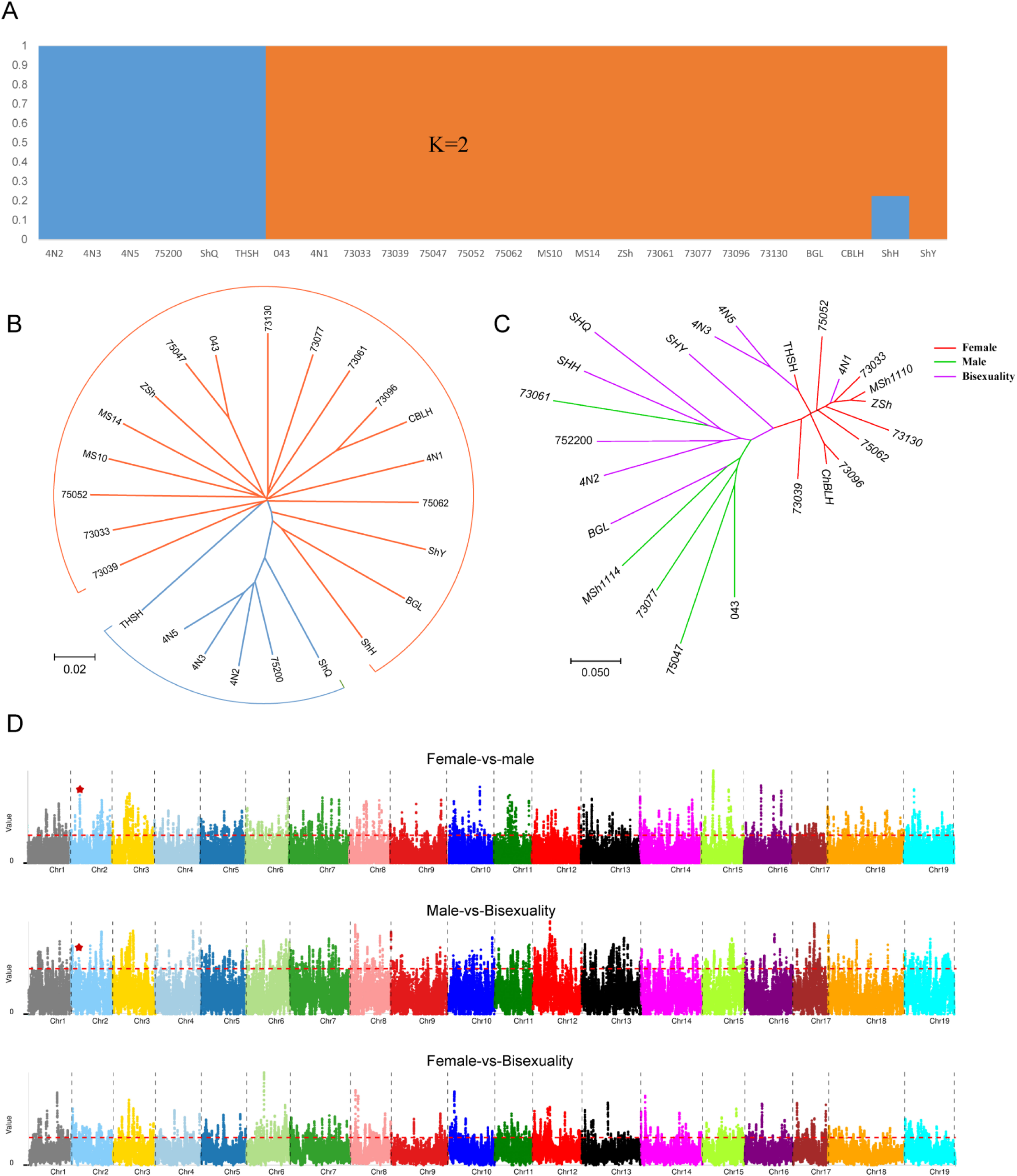
Population structure analysis and selective sweep analysis of 24 *Va* individuals. (A) Population structure of 24 *Vitis amurensis* individuals (There are some individuals whose names use abbreviations here. Please refer to table s14 for the corresponding relationship between abbreviations and full names of individual names). (B) An evolutionary tree based on the whole-genome SNPs of 24 *Va* individuals. (C) An evolutionary tree based on the SNPs on the SDRs of 24 *Va* individuals. (D) A Manhattan diagram of the selective region. The top subfigure shows the selective region from the comparison of female and male groups comparison. The middle subfigure is the selective region from the comparison of female and hermaphroditic groups. The bottom subfigure shows the selective region from the comparison of male and hermaphroditic groups. The red dashed line represents the threshold of θπ ratios and FST (top 5%). The blue asterisk represents the selective region which harbored SDR genes.

Therefore, a selective sweep analysis was performed according to the sex groups rather than the population structure. One of the selected regions inferred (both θπ ratios and FST in the top 5% range) from the comparison of male and female accessions totally overlapped the Amur grape putative SDR. We found that the Amur grape putative SDR contains 16 genes (Vitis02G0494, Vitis02G0495, Vitis02G0496, Vitis02G0497, Vitis02G0498, Vitis02G0499, Vitis02G0500, Vitis02G0501, Vitis02G0502, Vitis02G0503, Vitis02G0504, Vitis02G0505, Vitis02G0506, Vitis02G0507, Vitis02G0508 and Vitis02G0509), respectively homologous to *Vv* SDR genes associated with sex (Figure 6D). Another selected region inferred between male and hermaphroditic groups overlapped part of the Amur grape putative SDR gene Vitis02G0494 (Figure 6D). However, none of the selected regions inferred between female and hermaphroditic groups overlapped the Amur grape putative SDR (Figure 6D).

Previous studies showed that the SDR of *Vv* M and H haplotype genes include those respectively encoding PPR-containing proteins, a YABBY transcription factor (VvYABBY3), a VviSKU5 (Skewed5), a beta-fructofuranosidase, an aldolase, a trehalose-6-phosphate phosphatase (TPP), an inaperturate pollen1 (VviINP1), an exostosin family protein, KASIII, two TPR-containing protein, a PLATZ transcription factor, three Flavin-containing monooxygenases (FMO), a hypothetical protein (VviFSEX), a WRKY transcription factor, and an Adenine phosphoribosyltransferase (VviAPT3). The SDR of *Vv* F haplotypes included genes encoding a PPR-containing protein, a YABBY transcription factor (VviYABBY3), a VviSKU5, a beta-fructofuranosidase, an Aldolase, a Trehalose-6-phosphate phosphatase (TPP), an inaperturate pollen1 (VviINP1), an exostosin family protein, a KASIII, a PLATZ transcription factor. The F haplotypes also had four flavin-containing monooxygenases, a hypothetical protein (VviFSEX), a WRKY transcription factor, and an Adenine phosphoribosyltransferase (VviAPT3), respectively (Massonnet *et al*., 2020).

In *Va*, the F haplotypes included 16 genes encoding a PPR-containing protein, a YABBY transcription factor (VviYABBY3), a VviSKU5, a beta-fructofuranosidase, a fusion protein of aldolase and trehalose-6-phosphate phosphatase (TPP), an inaperturate pollen1 (VviINP1), an uncharacterized protein (homologous to VIT_202s0154g00120 of *Vv* PN40024), an exostosin family protein, a KASIII, and a PLATZ transcription factor. Besides, the F haplotypes contained three flavin-containing monooxygenases, a hypothetical protein (VviFSEX), a WRKY transcription factor, and an adenine phosphoribosyltransferase (VviAPT3), respectively. The SDR of *Va* F haplotypes had an additional uncharacterized protein gene (Vitis02G0500) and lost one FMO gene, as compared to the *Vv* Cabernet Sauvignon F haplotypes (Figure 7).

**Figure 7.** Genes of *Vv* and *Va* SDRs. Genes *of Vv* H, F hap *SDRs* are from *Vv* Cabernet Sauvignon. Genes of *Vv* M hap SDR are from *Vv. sylvestris*. Genes of Va F hap are from Zuoshan No.1. The gene with a arrowhead is the *Va* Uncharacterized protein gene.

In *Vv*, the female-specific DNA polymorphisms at -13, -5, and +2 bp may reduce transcription and/or alter mRNA decay of the female PLATZ transcription factor allele on *Vv* SDR (Iocco-Corena *et al*., 2021). However, *Va* lacks these DNA polymorphisms upstream of the ATG of the PLATZ transcription factor gene in the Amur grape putative SDR. In other words, there is no difference of DNA polymorphisms between female, hermaphrodite, and male *Va* individuals within 20 bp upstream of the PLATZ transcription factor gene.

We found some other selected regions may be related to sex-determination (θπ ratios and FST estimates in the top of 5%). For example, the region (chr3: 6730001bp-8840000 bp) from the comparison of male and female accessions, the region (chr16:3565001bp-4830000bp) from the comparison of male and female accessions, the region (chr16:15590001bp-18015000bp) from the comparison of male and female accessions, and the region (chr17:11450001bp-13785000bp) from the comparison of male and hermaphroditic accessions. In these regions, there are the MADS-box protein AGL62 genes (Vitis03G0709, Vitis03G0710, Vitis03G0711 and Vitis03G0712) from chr3: 6730001bp-8840000 bp. We found that there are the Pentatricopeptide repeat-containing protein genes in chr16:3565001bp-4830000bp (Vitis16G0214 and Vitis16G0215), in chr16:15590001bp-18015000bp (Vitis16G0545), and in chr17:11450001bp-13785000bp (Vitis17G0924). These Pentatricopeptide repeat-containing genes are the homologs of putative SDR gene Vitis02G0494 (a Pentatricopeptide repeat-containing gene).

## Discussion

*Vitis* mainly include the Eurasian grape, East Asian grape, and the American grape. The sequence of the *Vv* genome (PN 40024), of a Eurosian grape, was published in 2007 (Jaillon *et al*., 2007) and incurred researches to understand grape biology (Wang *et al*., 2018a; Wang *et al*., 2018b; Wang *et al*., 2019; Liang et al., 2019). Given that its genome has only undergone one polyploidization, the ECH, after split from the basal endicots, it is often used as outgroup reference for studying the other endicot genomes (Wang *et al*., 2018a). The sequence of *Vr* RGM genome, of an American grape, was published in 2019 (Girollet *et al*., 2019). The Cabernet Sauvignon genome sequence, a *Vv* variety, was later published, being assembled using the Hi-C technology (Massonnet *et al*., 2020).

Here, we assembled the genomes of *Vitis amurensis*. The above analysis showed that the assembled *Va* genome sequence reported here is much improved as to the previous one. Based on the present genome sequence, we discovered a likely assemble error in chromosome 13 of the previous grape genome sequence.

The availability of multiple grape genome sequences allows for comparing and analyzing gene sequences of several grape species so that we could detect the evolution history and other important comparative genomics information across the grapes. In the present study, gene synteny analysis showed the arrangement order of orthologous genes between *Va*, *Vv*, and *Vr* genomes are much conservative. We have to note here, genome instability due to the ECH, as ancient as 130 Mya, contributed to the divergence of the grape genomes. Actually, a characterization of homologous genes showed that *Vr* is the most conservative one among the three. *Vr* contains more paralogous genes produced by the ECH, comparing with *Va* and *Vv*, showing the conservativeness of the *Vv* genome, and implying that *Vr* may much resemble their common ancestor. Moreover, more orthologous genes were preserved between *Vr* and the other two grapes (*Vv* and *Va*) than between *Vv* and *Va*. Besides, phylogenetic analysis supported that *Vr* was the first one to split from the other grapes. These show that *Vr* genome may be taken as a better reference for *Vitis* biology and even all the eudicots. The present finding that a 130-Mya polyploidization contributed to the species divergence of grapes reminded us that it may has played a non-negligible role in grape divergence and genetic innovation, as to a similar finding in rice and other grasses (Wang *et al*., 2011). Possibly, the genomic feature of thousands of duplicated genes in crops could be manipulated to breed high-yield and/or high-quality crops.

Species-specific genomic segments also make up a large proportion of the genomes for all the three species, implying significant genomic differences among the three species. SV may generate phenotypic differences (Chawla *et al*., 2021), for example, *Va* being more cold-resistant than the other two species. Biased sequence preservation in *Va* may have contributed to its capability to resist coldness. Sequence variations, especially incurred by the ECH, may generate phenotypic differences. Here, we found that the *Va*-specific PAV genes contain cold resistance-related genes (CRG), such as those encoding the ethylene-responsive transcription factors and cold-regulated 413 inner membrane proteins. The robustness of CRG regulatory networks in *Va* is higher than those in *Vv* and *Vr*. This showed that the stronger cold resistance of *Va* may be related to the preferential preservation of cold-related genes, and *Va* cold-related genes may constitute a more robust and effective interaction network. A study of CRGs in angiosperms, including *Vv*, supported the expansion of them was significantly related to the occurrence of polyploidization (Song *et al*., 2020). The present study provides further evidence that the ECH, though occurred 130-150 million ago, still contribute to enhance resistance to coldness in plants.

Multi-omics analysis of the *Va* was performed to understand the mechanism underlying the regulation of nutrient accumulation in the *Vitis amurensis* over its developmental period. *Va* is sourer than *Vv*, a factor that needs urgent resolution by breeders, in that high sourness makes the fruits taste unpleasant, and the wine tastes unbalanced. The present analysis revealed that the total sugar increased continuously during the “Zuoshan No.1” berry development, while the total sugar content in this species/variety was almost equal to that in *Vv*. Further analyses revealed that the total acid content decreased significantly during “Zuoshan No.1” berry development. Wide target metabolome analysis showed no decrease in tartaric acid content during berry of “Zuoshan No.1” development, while those of citric acid and several malic acid types decreased significantly in “Zuoshan No.1”. Combined transcriptome and metabolome analyses revealed that sucrose-6-phosphatase, alpha, alph-trehalase, and beta fructofuranosidase genes jointly regulate the accumulation of sucrose and trehalose during “Zuoshan No.1” development. The DEGs during S1-S4 comprises the ripening-related protein-like gene and chalcone synthase gene, which is the first enzyme activated the flavonoid biosynthetic pathway (Waki *et al*., 2020), UDP-glucose: flavonoid 3-O-glucosyltransferase gene, which promotes anthocyanin accumulation in grape (Yamazaki *et al*., 2021), SWEET gene, which is a sugar transporter and uniporter gene (Chong *et al*., 2014), ethylene-responsive transcription factor 3-like gene, certaining bHLH, MYB transcription factor genes, and WD repeat-containing protein genes. Most of these genes may be involved in *Va* berry ripening. Therefore, we infer that these DEGs may play a central role in the process of *Va* berry ripening.

The *Va* genome was further analyzed to understand the relationship between different *Vitis amurensis* phenotypes and genotypes. Given the scarcity of hermaphroditic *Vitis amurensis* in nature, we explored the sex determination region in this grape. Previous studies have uncovered the SDR of *Vv* (Massonnet *et al*., 2020). The three *Vv* types (varieties) are determined by the genotype at the SDR. Males are heterozygous for the male and female haplotypes (MF), females are homozygous (FF), and cultivated *Vv* hermaphrodites are either homozygous for hermaphrodite haplotypes (HH) or heterozygous (HF). Previous studies have shown that the SDR of *Vv* M and H haplotypes have 2 TPR-containing protein genes and 3 Flavin-containing monooxygenases (FMO), and the SDR of *Vv* F haplotypes contain 4 Flavin-containing monooxygenases (FMO) without TPR-containing protein gene (Massonnet *et al*., 2020). This implies that the type or number of genes in the SDR of grapes vary between sexes.

Further analyses to clarify whether SDR is also present in *Va* revealed that one fragment from *Va* is homologous to *Vv* SDR in *Vitis amurensis* and call it the Amur grape putative SDR. In the present study, we sequenced the female *Va* genome. Therefore, further analyses were performed to explore whether the types and numbers of genes in female *Va* putative SDR were consistent with those in the female *Vv* SDR. Selective sweep analysis of male, female, and hermaphroditic individuals revealed one selective region from the comparison of male and female accessions overlapped total the Amur grape putative SDR including 16 genes all homologous to *Vv* SDR genes (such as APT3 gene, PLATZ transcription factor gene and TPP gene). A previous study inferred that VaAPT3 gene (orthologous to Vitis02G0508 in the present *Va* genome), located in Amur grape putative SDR, was associated to sex determination (Men *et al*., 2021). Recently, the PLATZ transcription factor gene has been found to determine the grape’s sexuality (Iocco-Corena *et al*., 2021). The TPP gene also located in Amur grape putative SDR and TPP genes in many species have also been associated with sex determination, flower development, reproduction, and synthesis of plant hormones, among other functions (Kataya *et al*., 2020; Qiu *et al*., 2020). PtTPPs displayed a specific expression pattern in seven developmental stages of Populus male and female floral buds (Gao *et al*., 2021). Trehalose-6-phosphate phosphatase (TPP) controls inflorescence architecture in maize through sugar signal modification (Satoh-Nagasawa *et al*., 2006). This suggests that the hypothetical *Va* SDR may be related to sex determination. One selected region was from the comparison of male and hermaphroditic accessions overlapped part of Amur grape putative SDR including only one gene (PPR-containing gene), which is homologous to the *Vv* SDR gene PPR-containing gene. Additionally, some selective regions may be related to *Va* sex determining. Previous study indicated that some PPR-containing proteins genes could restore fertility to cytoplasmic male-sterile plants (Bentolila *et al*., 2002; Koizuka *et al*., 2003; Kazama *et al*., 2003; Hu *et al*., 2012). A study on RNA-Seq analysis of three flower sex types in grapevine showed some grape PPR-containing genes could be essential for carpel development (Ramos *et al*., 2014). The study also inferred that PPR-containing gene may be essential for the perfect development of sexual floral organs. Here, we found that some other selected regions comprises many PPR-containing protein genes, allowing for further exploration of their likely contribution to sex determination.

However, no selected region overlapped the Amur grape putative SDR from the comparison of male and hermaphroditic accessions was found. We found that the type and arrangement order of all genes in the putative SDR of female *Va* and the SDR of *Vitis vinifera* were similar. However, compared with *Vv* F haplotypes, the putative SDR of female *Vitis amurensis* has lost one FMO gene and contains one extra Uncharacterized protein. The Uncharacterized protein gene has never been found in SDR of *Vv* Cabernet Sauvignon and *Vv. sylvestris* (Massonnet *et al*., 2020). The role of the Uncharacterized protein gene in *Vitis amurensis*sex determination needs further exploration.

In the other selective region (chr17:11450001bp-13785000bp), we found from the comparison of *Va* male and hermaphroditic accessions, there are many MADS-box protein AGL62 genes. Previous study inferred MADS-box protein genes may be also related to developing male and female flowers (Hardenack *et al*., 1994; Ainsworth *et al*., 1995). We consider that the region (chr17:11450001bp-13785000bp) may be also related to *Va* sex-determining. Plus those selective regions that have been found to contain the PPR-containing proteins genes, it should be explored whether *Vitis amurensis* sex may be determined by several different genomic regions in the future.

## Materials and methods

### Plant materials and DNA sequencing

Fresh leaves and stems of *Vitis amurensis* cv. “Zuoshan No. 1” were sampled for DNA extraction and sequencing. Total genomic DNA was extracted using the CTAB (cetyltrimethylammonium Ammonium Bromide)) method (Tel-Zur *et al*., 1999). The library for ONT sequencing (Oxford Nanopore Technology, Oxford, UK) was constructed using large (>15 kb) DNA fragments with the SQK-LSK109 Ligation Sequencing Kit and sequenced using the ONT platform. Adapters and low-quality nucleotides (with <7 mean quality score) were trimmed off. Paired-end libraries with 350 bp insert sizes were constructed following the manufacturer’s protocols and sequenced using the MGIseq 2000 platform (MGI Tech Co. LTD, Guangdong, China). The MGIseq reads were filtered using the SOAPnuke1.5.6 online software (https://github.com/BGI-flexlab/SOAPnuke). For the high-throughput chromosome conformation capture (Hi-C) analysis, fresh leaves and stems of *Vitis amurensis* cv. “Zuoshan No. 1” were treated following previous methods (Yang *et al*., 2020).

### Genome size and heterozygosity estimation

The genome size was estimated from the MGIseq reads using k-mer analysis, and the k-mer depth-frequency distribution was generated using the jellyfish software (Marçais and Kingsford, 2011). The genome size and heterozygosity were calculated using the genomeScope software (Vurture *et al*., 2017).

### *De novo* genome assembly

Long ONT reads were corrected using Canu (https://github.com/marbl/canu/releases), and *de novo* assembled using the Smartdenovo software (https://github.com/ruanjue/smartdenovo). The Racon (https://github.com/isovic/racon) and Medaka (https://github.com/nanoporetech/medaka) software were applied to polish the assembled contigs. The polished, assembled data were then corrected using Pilon (v.1.22, https://github.com/broadinstitute/pilon/). Next, the Haplomerger2 pipeline reduced the assembly to ∼522 Mb. After that, the purged assembled contigs were anchored into 19 pseudochromosomes using the Juicer with the parameters “-g draft -s MboI” and 3D de novo assembly (3D-DNA) pipeline with the parameters “-m haploid -r 2” (Durand *et al*., 2016; Dudchenko *et al*., 2017).

### Genome annotation and gene prediction

Transposable elements (TEs) were predicted using combined homology-based comparisons with RepeatMaskerv4.0.7, RepeatProteinMask v4.0.7, and *de novo* approaches with Piler (Edgar and Myers, 2005) (http://www.drive5.com/piler/), RepeatScout, and RepeatModeler. Tandem repeats were identified using the Tandem Repeats Finder v4.09 (Benson., 1999) (http://tandem.bu.edu/trf/trf.html) and LTR_FINDER v1.06 (Zhao and Hao, 2007) (http://tlife.fudan.edu.cn/ltr_finder/). Furthermore, the *Va* protein-coding gene set was deduced by *de novo*, homology, and evidence-based gene prediction (transcriptome data) (Guo *et al*., 2021). The transcript evidence included transcripts assembled from the RNA-Seq data of different tissues (leaf, stem, flowers, tendril, and fruit; Table S9). Next, the predicted genes were functionally annotated using a previous method (Guo *et al*., 2021). Moreover, tRNA was identified using tRNAscan-SE 1.3.1 (Lowe and Eddy, 1997) (http://lowelab.ucsc.edu/tRNAscan-SE/), BLASTN identifiedrRNA, while INFERNAL (http://infernal.janelia.org/) identified miRNA and snRNA.

### Genome homology inference

Protein sequences from one plant were searched against themself and those another plant genome using BLASTP (Camacho *et al*., 2009) to find the best, secondarily best, and the other matches with E_value<1E-5. Dot plots were produced using the WGDI package of Python (Sun *et al*., 2021). Colinear genes were inferred using the -icl subprogram contained in WGDI package with default parameters. Nucleotide substitution rates were estimated between colinear homologous genes using the YN00 program in the PAML (v4.9h) package implementing the Nei-Gojobori approach (Yang, 2007).

### Species tree

A rooted species tree was inferred using the Orthofinder (version 2.5.4) with the “-M msa” parameter (Emms and Kelly, 2015; Emms and Kelly, 2019) based on 5929 single-copy genes.

### Identification and characterization of cold-related genes

The cold-related genes in the three grape species and gene network robustness were determined following previous methods (Song *et al*., 2020).

### Structural variations analysis

The SV analysis was performed using the NUCmer program embedded in MUMmer with the parameters “-mumreference -g 1000 -c 90 -l 40”. (Yu *et al*., 2021). Genomic-specific segment analysis was performed as follows. First, the query genome sequences was split into 500bp windows with a overlapping step size of 100bp. and then all the 500bp windows subsequnces was aligned against the references genome by BWA-mem with the parameters “-w 500 -M”. The 500bp window that failed to align or aligned with less than 25% coverage were defined as the genomic-specific segments. The genes within the genomic-specific segments, were defined as the specific-genes of the query genome (species-specific PAV genes). Structural variation plots was drawn using ggplot2.

### Widely targeted metabolomic analysis

Berries from four growth stages were used for this study, including Stage1 (S1, the late period of berry expansion), Stage 2 (S2, Veraison), Stage 3 (S3, the period when berries change color completely), and Stage 4 (S4, maturity stage). Three biological replicates from each stage were used for metabolome analysis. The Metware Biotechnology Co., Ltd. (Wuhan, China) performed the metabolome extraction and analysis using previous methods (Ma *et al*., 2021). The metabolites were identified using the Metware database (MWDB), and metabolite abundances were determined according to the metabolite peak areas. Metabolites were considered as differentially accumulated when the variable importance in projection (VIP) was ≥1, and the absolute log2 (fold change) was ≥1. Metabolic pathways were constructed according to the KEGG database.

### Total sugar and titratable acidity content analysis

Berries from four growth stages (S1, S2, S3 and S4) were grind into homogenate used for this study. Three biological replicates from each stage were used for metabolome analysis. We analyzed the total sugar content using sulfuric acid-anthrone colorimetric method and the instrument used is Lanbda 365 ultraviolet/visible spectrophotometer. We analyzed the titratable acidity content using acid-base titration and the standard solution is Sodium hydroxide standard solution (0.05 mol/L).

### RNA-seq

Just as we analyzed the metabolome, berries from four stages, S1, S2, S3, and S4, were used for the RNA-seq analysis. Three biological replicates from each berry stage were used for RNA-seq analysis. Novogene Co. LTD performed the RNA-seq experiments on the Illumina Novaseq 6000 platform (Illumina, CA, USA) following the manufacturer’s instructions and previously described steps (Zhang *et al*., 2021). All paired-end reads were mapped to the *Va* genome using Hisat2 v2.0.5. Expression levels were calculated using the fragments per kilobase of exon model per million mapped fragments (FPKM). The DEGSeq R package (1.20.0) was used to identify differentially expressed genes (DEGs). Genes with adjusted *P* value < 0.05 and fold changes ≥ 2 or ≤ -2 were identified as differentially expressed genes (DEGs).

### Integrated analysis of metabolome and RNA-seq

For the combined analysis of all metabolome and transcriptome, the canonical correlation analysis (CCA) was performed on the metabolome and RNA-seq data using CCA package in the R statistical environment (Gonzalez *et al*., 2008). Next, the WGCNA (Weighted correlation network analysis) was performed on all RNA-seq and metabolome data using the WGCNA package in the R statistical environment.

### Population genetic analysis

Essentially, 24 Amur grape genotypes were re-sequenced based on the *Va* genome assembled in this studyand used for calling SNPs. Concurrently, the ANNOVAR package was employed for population genetic analysis. Ten folds *Va* genome size of clean reads data of 24 kind of Amur grapewere separately obtained on the MGI2000 platform. The SNPs in LD were filtered using PLINK with a window size of 50 SNPs (advancing 5 SNPs at a time) and a 0.5 r^2^ threshold. The PCA was conducted using GCTA (v1.25.2) (Yang *et al*., 2011). Furthermore, the population structure was analyzed using FRAPPE, and the MLtree was constructed using SNPhylo (Lee *et al*., 2014) to clarify the phylogenetic relationships.

### Selective sweep analysis

Selective sweep analysis was performed using a previous method (Lin *et al*., 2022). In briefly, Fst and θπ were used to detect candidate selective regions between the three grape populations, Fst and θπ were calculated using PopGenome (Brigida *et al*., 2016; Pfeiferet *et al*., 2014); regions with both θπ ratios and FST estimates in the top of 5% were considered as the selection regions.

### Accession number

The *V*. *amurensis* genome project was deposited at NCBI under BioProject number PRJNA868106 and BioSample SAMN30308461.

## Author Contributions

P.W., Y.Z., Y.Y., H.Z. and B.L. designed the experiments. Q.M., T.D., H.L., F.W., S.F. and Q.Z. performed the experiments and wrote the manuscript. F.M., A.L., Z. M. and T. Z. analyzed the data. J.J., Y.Z. and X.W. edited the manuscript.

## One Sentence Summary

We presented a high-quality *Vitis amurensis* genome sequence and conducted an in-depth study of grape genome variation, sex determination, and cold resistance, contributing to understanding of grape biology and breeding fine grapes.

The author responsible for distribution of materials integral to the findings presented in this article in accordance with the policy described in the Instructions for Authors (https://academic.oup.com/pphys/pages/General-Instructions) is Xiyin Wang.

## Acknowledgements

This research was supported by the Key Research and Development Project of Shandong Province (2021LZGC025; 2022CXGC010605), Improved Variety Program of Shandong Province (2020LZGC008), Shandong Natural Science Funding (ZR2020QC145), Scientific Research Guide Foundation of Shandong Academy of Grape (SDAG2021A01; SDAG2021A02), Shandong Academy of Agricultural Sciences, Introduction and Training of High-level Talents (CXGC2022E15), National Horticulture Germplasm Resources Center-Amur Grapevine Germplasm Resources Branch Center (NHGRC2022-NH04) and Species and variety Germplasm resources protection project (2130135).

## Supplementary tables

**Table S1 Statistics of gene annotations for *Va* Zuoshan No.1 genome.**

**Table S2 Number of Homologous Genes Residing in Blocks within each Genome or between Genomes.**

**Table S3 Kernel Function Analysis of *K_s_* Distribution Related to Duplication Events Within each Genome and Between Genomes.**

**Table S4 Statistics of PAV regions between *Va* genome and *Vv* genome or *Va* genome and *Vr* genome.**

**Table S5 *Va-*specific PAV genes between *Va* genome and *Vv* genome.**

**Table S6 *Vv-*specific PAV genes between *Va* genome and *Vv* genome.**

**Table S7 *Va-*specific PAV genes between *Va* genome and *Vr* genome.**

**Table S8 *Vr-*specific PAV genes between *Va* genome and *Vr* genome.**

**Table S9 Information of 24 *Vitis amurensis* individuals.**

## Supplementary figure legends

**Figure S1. Dotplot of homologous genes between *Va* Zuoshan No.1 and *Va* IBCAS1988 genomes**

**Figure S2. The evolutionary tree based on the 3 grapes and Arabidopsis or Boxwood.**

**Figure S3. Structural Variation analysis plot based on ggplot2.**

**Figure S4. Tree of NBS family genes of the three species.**

**Figure S5. The distribution of NBS family genes on the chromosomes of the three species. Lines represent Ks values of NBS gene pairs that are less than 0.025 (red), larger than 0.025 but less than 1 (orange), and larger than 1 but less than 1.5 (grey).**

**Figure S6. The total sugar and titratable acid content of *Vitis amurensis* fruit at different stages.**

